# A quantitative approach to measure and predict microbiome response to antibiotics

**DOI:** 10.1101/2023.01.27.525904

**Authors:** Vince Tu, Yue Ren, Ceylan Tanes, Sagori Mukhopadhyay, Scott G. Daniel, Hongzhe Li, Kyle Bittinger

**Affiliations:** Division of Gastroenterology, Hepatology, and Nutrition, Children’s Hospital of Philadelphia, Philadelphia, Pennsylvania, United States; Center for Clinical Epidemiology and Biostatistics, Perelman School of Medicine, University of Pennsylvania, Philadelphia, Pennsylvania, United States; Division of Neonatology, Children’s Hospital of Philadelphia, Philadelphia, Pennsylvania, United States; Department of Pediatrics, University of Pennsylvania Perelman School of Medicine, Philadelphia, Pennsylvania, United States; Center for Pediatric Clinical Effectiveness, Children’s Hospital of Philadelphia, Philadelphia, Pennsylvania, United States

## Abstract

**Background:** Although antibiotics induce sizable perturbations in the human microbiome, we lack a systematic and quantitative method to measure and predict the microbiome’s response to specific antibiotics. Here, we introduce such a method, which takes the form of a Microbiome Response Index (MiRIx) for each antibiotic. Antibiotic-specific MiRIx values quantify the overall susceptibility of the microbiota to an antibiotic, based on databases of bacterial phenotypes and published data on intrinsic antibiotic susceptibility.

**Results:** We applied our approach to five published microbiome studies that carried out antibiotic interventions with vancomycin, metronidazole, ciprofloxacin, amoxicillin, and doxycycline. We show how MiRIx can be used in conjunction with existing microbiome analytical approaches to gain a deeper understanding of the microbiome response to antibiotics. Finally, we generate antibiotic response predictions for the oral, skin, and gut microbiome in healthy humans.

**Conclusions:** Our approach is implemented as open-source software and is readily applied to microbiome data sets generated by 16S rRNA marker gene sequencing or shotgun metagenomics.

## Background

Data generated by human microbiome research has led to insights into the composition of microbial communities, including their functions, interactions, metabolism, and role in host health. Antibiotic use is a leading factor affecting microbiome composition in humans^1,2^, altering the microbiome directly^3,4^ and modulating the effects of diet^5^ and disease^6^ on the microbiome. Research has shown that taking antibiotics even for a short duration can have long-lasting consequences for the host microbiome, leading to a decrease in taxonomic diversity^7,8^. Despite the potency of antibiotics as influencers of the microbiome, and the wealth of information available on bacterial response to antibiotics, we lack the means to systematically apply this knowledge in the analysis of microbiome data.

Researchers have previously developed several approaches to quantify the complex interactions of microorganisms with antibiotics, outside of microbiome data analysis. The Drug Resistance Index (DRI) is a measurement of the overall level of microbial resistance of a single pathogen and captures the relationship between antibiotics susceptibility and antibiotic use from a temporal and spatial standpoint into a single indicator^9,10^. The drug effectiveness index (DEI) is an adaptation of the DRI for multiple select bacterial species and combines the probability of a certain species causing infection with the relative frequency of prescribing an antibiotic and the rate of resistance of a microorganism to that antibiotic^11^. The Multiple Antibiotic Resistance Index (MAR Index) was developed to measure the rate of multiple drug resistance of the isolates and the total number of tested antibiotics^12^. Other indices measuring antibiotic resistance take into account the availability of the drug and the proportion of bacteria associated with the infection^13^. For marine and freshwater environments, an index that quantifies antibiotic resistance was developed using functional metagenomics and sequence similarity to previously characterized antibiotic resistance genes^14^. Machine learning and data mining techniques have also been applied to calculating various versions of the DRI^15^.

The existing indices summarize the effectiveness of an antibiotic for important pathogens, but none are comprehensive enough to estimate the impact of antibiotics on a complex community of commensal along with potentially pathogenic bacteria. Thus, we were motivated to craft an approach to quantify and predict the anticipated effects of antibiotics on bacterial communities as they exist in the human body.

Here, we introduce the concept of an antibiotic-specific microbiome response index, which is based on the ratio of susceptible to non-susceptible organisms in a microbial community. The proportions of susceptible and non-susceptible organisms are computed based on the taxonomic composition of the microbiota within a sample. The index can be used to quantify the degree to which the balance of organisms in a community is tilted towards non-susceptible (hereafter, resistant) organisms. Recognizing that a complex bacterial community is likely to respond in unanticipated ways due to ecological factors, our approach allows the off-target effects of an antibiotic to be assessed. Moving beyond simple measurement, our approach can be used to *predict* the microbiome’s response to different antibiotics. Furthermore, a microbiome response index can be combined with microbiome diversity or taxonomic abundance to enrich and expand data analysis. Here, we describe our new approach and show how it can be used to further human microbiome research.

## Methods

Five previously published data sets were downloaded and used to evaluate the approach presented here.

### Data acquisition and processing

For the studies with shotgun metagenomic sequence data^16,17^, FASTQ files were downloaded from the NCBI Sequence Read Archive (SRA). Bioinformatics processing was carried out using the Sunbeam metagenomics pipeline^18^ using default parameters. Briefly, sequence reads were trimmed to remove adapters and low-quality sequence with cutadapt^19^ and Trimmomatic^20^, respectively. Reads aligning to the human genome or that of phage ϕX174 were removed by alignment with BWA^21^, and low-complexity sequences were removed with komplexity^18^. Taxonomic assignments were generated with Kraken^22^.

For the study from Sprockett et al.^23^, FASTQ files were downloaded from the SRA. For the study from Boynton et al.^24^, FASTQ files were obtained directly from the authors. Bioinformatics processing was carried out using the QIIME2 pipeline^25^. Sequences were trimmed to 263 bp based on sequence quality prior to denoising with DADA2^26^. Taxonomic assignments were generated using the built-in naïve Bayes classifier in QIIME2^27^ (q2-feature-classifier).

To access the Human Microbiome Project (HMP) dataset, we used the R package HMP16SData to download processed data and taxonomic assignments^28^.

Microbiome response index values were calculated with the mirix R package, presented here, using the built-in databases for taxon phenotypes and antibiotic-specific susceptibility.

### Statistical analysis

For studies without repeated measurements (Basolo et al., Cabral et al., Human Microbiome Project), we used linear models to compare values of the microbiome response index, taxonomic abundance, Shannon diversity, and level of dysbiosis between sample groups. For studies with repeated measurements of the same subject (Sprockett et al., Willmann et al., Boynton et al.), we used linear mixed effects models and included a random intercept for each subject. The relative abundances of bacterial taxa were log-transformed before comparison. P-values for taxonomic comparisons were adjusted to control for the false discovery rate using the method of Benjamini and Hochberg^29^. We used Pearson correlation to analyze correlations between the microbiome response index and Shannon diversity or dysbiosis and carried out a test of correlation to obtain p-values. We used the PERMANOVA test to compare microbiome community composition between groups^30^.

## Results

### A quantitative approach to measure microbiome response to antibiotics

The Microbiome Response Index (MiRIx) summarizes the overall susceptibility of the microbial community in a microbiome sample to a specific antibiotic. The index is computed as the log of susceptible organisms over resistant organisms. Communities with a positive index harbor mostly susceptible organisms and would be expected to respond strongly to the introduction of antibiotics. Communities with a negative index harbor mostly resistant organisms and would be expected to respond less. Antibiotic intervention is expected to drive the index down, eliminating susceptible organisms and thus making the community less responsive to further exposure to the antibiotic.

In Figure 1A, we show three examples to illustrate how the index works. In Sample A, 80% of the bacterial community is susceptible to the antibiotic. The MiRIx value for Sample A is positive, log(80 / 20) = 0.6, indicating a majority of susceptible organisms and a potential for the community to respond vigorously to antibiotic intervention. Sample B is balanced slightly in favor of susceptible organisms. Thus, the value of the index is close to zero but slightly positive. In Sample C, the microbiota is dominated by resistant organisms, with susceptible organisms accounting for only 10% of the total abundance. Consequently, the MiRIx value for Sample C is negative, at log(10 / 90) = -0.95.

**Figure 1.**
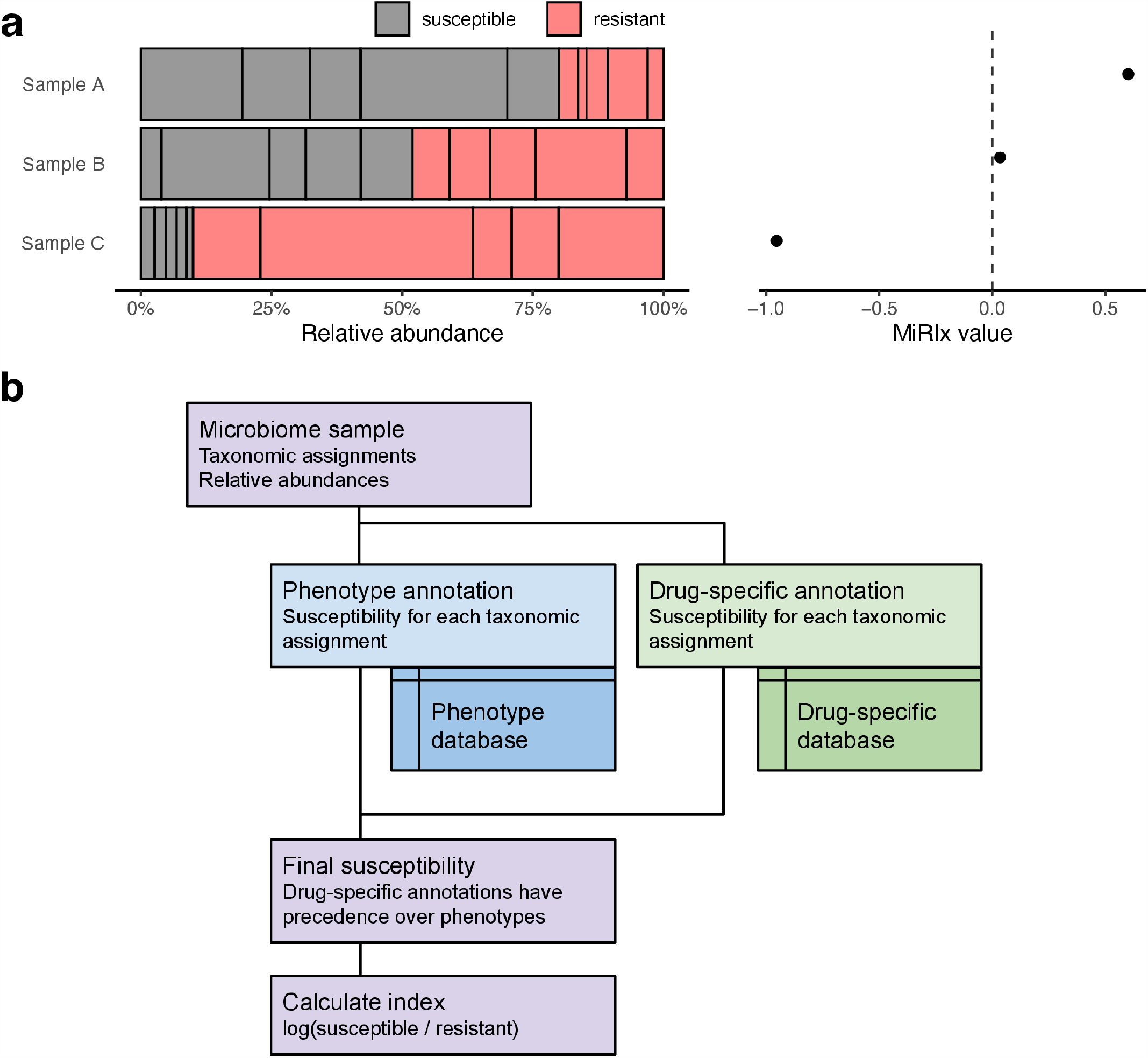
Diagram of an antibiotic-specific microbiome response index (MiRIx). (a) Examples of three bacterial communities, showing susceptible and resistant taxa, alongside the value of the index. (b) Algorithm for computing values of the index. The inputs are taxonomic assignments and relative abundances for a microbiome sample. Using separate phenotype and drug-specific databases, taxa are annotated as susceptible or resistant. Finally, the index is computed from the ratio of susceptible to resistant bacteria.

To determine the susceptibility or resistance of a bacteria in the sample microbiome community, we rely heavily on databases of bacterial phenotypes and antibiotic susceptibility that we assembled from the scientific literature. Figure 1B gives an overview of how the index is computed. The input is taxonomic assignments and their relative abundances for a microbiome sample. For each taxonomic assignment, the phenotype database is searched to determine susceptibility based on phenotype. As an example, using the vancomycin index, we annotate Gram-positive bacteria as susceptible and Gram-negative bacteria as resistant. After annotating based on phenotype, a second database of drug-specific information is searched for additional annotations. For example, *Lactobacillus* is annotated as resistant to vancomycin, due to the inherent resistance of its Gram-positive cell wall. Next, the phenotype and drug-specific susceptibilities are merged, with the drug-specific susceptibility taking precedence. Taxa that are not represented in either database are not labeled as either resistant or susceptible. As a consequence, they do not enter into the computation. Finally, the value of the index is calculated by dividing the relative abundance of susceptible bacteria by the relative abundance of resistant bacteria and taking the base-10 logarithm.

The microbiome response index provides a summary measurement of the microbiome that can be used in statistical models to quantify the microbiome response to an antibiotic. In this way, the index is able to provide insight into the relative effects of disease and antibiotics. The index can highlight taxa that were predicted to be resistant to antibiotic intervention but were nevertheless observed to change in abundance. Moreover, the approach can be used to predict the degree to which different antibiotics will perturb the microbiome.

### Application of antibiotic-specific microbiome response indexes

We downloaded three publicly available microbiome data sets to demonstrate an analysis of microbiome response index in antibiotic intervention studies. The studies used vancomycin, metronidazole, and ciprofloxacin, respectively, and carried out either 16S rRNA marker gene sequencing or shotgun metagenomic sequencing.

In the first study, Basolo et al. treated ten adults with vancomycin and eight with placebo (Figure 2A)^16^. The fecal microbiome was collected following treatment and subjected to shotgun metagenomics. In the placebo group the vancomycin-MiRIx had a mean value of 0.016, corresponding to a roughly 50:50 mixture of bacteria annotated as susceptible and resistant. The mean value of the vancomycin-MiRIx was -0.91 in the vancomycin-treated group, indicating an increase in the ratio of resistant-to-susceptible bacteria (P=0.02).

**Figure 2.**
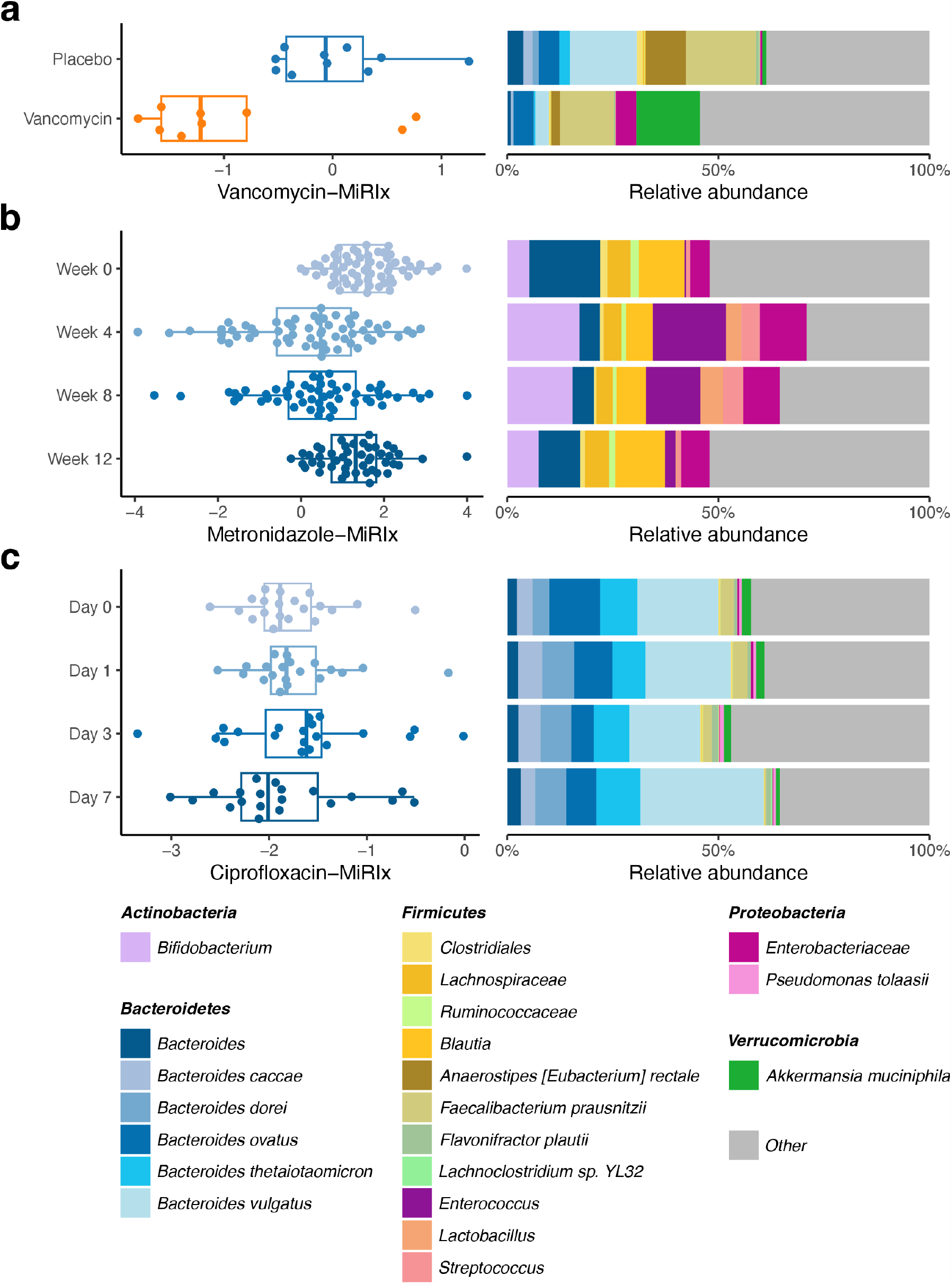
MiRIx applied to previously published studies. Observed MiRIx values and taxonomic abundances for three previously published human gut microbiome studies: (a) vancomycin (n=10) vs. placebo (n=8) from Basolo et al., (b) a metronidazole time course (n=67) from Sprockett et al., and (c) a ciprofloxacin time course (n=20) from Willmann et al.

An analysis of taxonomic abundance revealed differences that were generally in line with the susceptibility predicted from our databases (Additional file 1). Taxa that were more abundant in the vancomycin group included *Klebsiella oxytoca* (P=0.002), which was annotated as resistant due to its Gram-negative phenotype. *Lactobacillus rhamnosus*, annotated as resistant to vancomycin in the drug-specific database, was also increased (P=0.004) in the vancomycin group. Taxa that were found to be lower in abundance for the vancomycin group were largely Gram-positive, and thus were annotated to be susceptible. In contrast, the genus *Bacteroides* decreased in abundance for the vancomycin group (P=0.006) despite being annotated as resistant. Although *Bacteroides* has been observed to decrease in abundance with vancomycin treatment in other studies^31^, the association is not predicted based on the direct action of vancomycin on bacteria classified as *Bacteroides*. This suggests that the difference in *Bacteroides* is an ecological consequence of vancomycin’s effect on other community members. Such a result should rightly be highlighted in an analysis of this study.

In the second study, from Sprockett et al., the fecal microbiome of 67 adults was sampled longitudinally at 0, 4, 8, and 12 weeks, and metronidazole was administered from weeks 4 to 8 (Figure 2B)^23^. As expected, the value of the metronidazole-MiRIx was lower during the period where metronidazole was administered, relative to week 0 (P=10^−8^). At week 12, the metronidazole-MiRIx returned to its baseline value (P=0.24). Although the overall microbiome susceptibility to metronidazole returned to its baseline value, the taxonomic configuration remained different at week 12 relative to baseline (PERMANOVA test, P=0.001), indicating that the microbiome assumed a different state as susceptible bacteria increased during the post-antibiotic period. This interpretation was, in turn, supported by an analysis of relative abundance for metronidazole-susceptible and resistant taxa (Additional file 2).

In the third study, from Willmann et al., we selected a set of 20 human subjects who were treated with ciprofloxacin over a period of 7 days (Figure 2C)^17^. Fecal microbiome samples were collected before treatment on day 0, and then on days 1, 3 and 7 following treatment. Applying the ciprofloxacin-MiRIx to shotgun metagenomic sequencing data from the study, we observed that the value of the ciprofloxacin-MiRIx did not change between pre- and post-treatment days 1, 3 and 7 (P=0.3, 0.5, and 0.7, respectively). Although our analysis indicated no difference in the overall ratio of annotated susceptible-to-resistant bacteria, we noted differences in relative abundance for bacterial taxa during the treatment period. On days 3 and 7, our analysis indicated a decrease in abundance of 13 taxa annotated as susceptible, but with a compensating increase among the *Bacteroides*, which were also annotated as susceptible (Additional file 3).

Our observation of diverging responses among bacteria that we expected to be susceptible allowed us to follow up with a targeted analysis of antibiotic resistance genes. We did not detect an increase in genes conferring resistance specific to fluoroquinolones over the time course; rather, we observed a small decrease in several efflux pump genes on day 7 compared to day 0 (Additional file 4). We uncovered no evidence that the ciprofloxacin intervention increased the abundance of organisms with relevant antibiotic resistance genes. Thus, annotation of susceptible and resistant taxa enabled the analysis of metagenomic data to proceed along a more biologically relevant direction, even though the antibiotic intervention did not perturb the microbiome as expected.

### Microbiome response indexes for broad-spectrum antibiotics

Broad-spectrum antibiotics such as amoxicillin are widely used and of considerable interest to researchers studying the microbiome. However, they pose a challenge for our approach, which relies on drawing a contrast between susceptible and non-susceptible taxa. Here, we describe how we accommodate broad-spectrum antibiotics with our approach and demonstrate the results using previously published data sets for amoxicillin and doxycycline.

To compute MiRIx values for broad-spectrum antibiotics, we initially assume that all bacteria are susceptible to the drug. As with other antibiotics, we then apply a second round of annotations from a drug-specific database, which allows some taxa to be flagged as resistant if the literature indicates that resistance has been predominantly acquired. For example, *tet* genes that confer resistance to most tetracyclines can be found in 80% of *Bacteroides* species ^32^, and consequently *Bacteroides* is annotated as resistant to tetracyclines in our database. Following annotation of resistant taxa from the drug-specific database, the calculation proceeds as before. In short, the drug-specific annotation database takes on an increased level of importance for broad-spectrum antibiotics.

Cabral et al. profiled the effects of amoxicillin on the structure of the murine microbiome^33^. They collected fecal microbiome samples from 4 mice treated with amoxicillin and 4 mice with placebo, then carried out shotgun metagenomic sequencing. As expected, values of the amoxicillin-MiRIx were lower in the amoxicillin group relative to the control group (P=0.0003, Figure 3A). Correspondingly, the amoxicillin treatment group had increased relative abundance of several *Bacteroides* species such as *B. thetaiotaomicron, B. ovatus*, and *B. fragilis* (Additional file 5), which were annotated as resistant to amoxicillin and other penicillin-like compounds in our drug-specific database, based on previous literature^34^.

**Figure 3.**
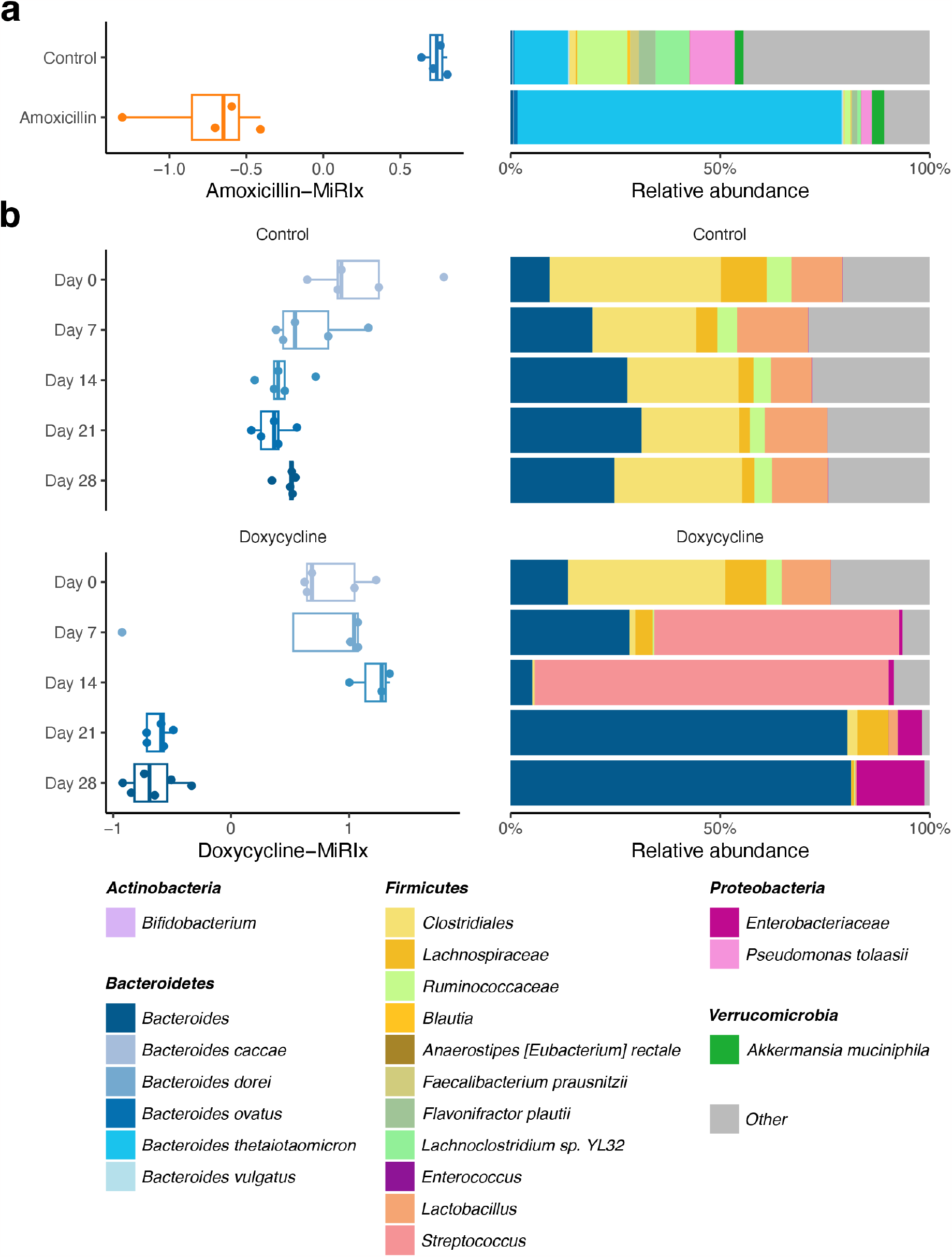
MiRIx applied to previously published studies of broad-spectrum antibiotics. Observed MiRIx values and taxonomic abundances for two previously published murine gut microbiome studies: (a) amoxicillin (n=4) vs. control (n=4) from Cabral and (b) a doxycycline (n=5) vs. control (n=5) time course from Boynton et al.

To evaluate the effectiveness of our method on another broad-spectrum antibiotic, we selected a study from Boynton et al., where mice were randomized into doxycycline treatment (N=5) and control groups (N=5)^24^. Fecal samples were collected at baseline and on days 7, 14, 21, and 28. The microbiota was profiled with 16S rRNA marker gene sequencing. In the treatment group, the doxycycline-MiRIx did not differ from baseline on days 7 and 14 (P=0.3) but decreased dramatically on days 21 and 28 (P=6×10^−5^, Figure 3B). Once again, the microbiome response index yielded additional insight to our analysis of taxonomic abundance: on days 7 and 14, communities in the treatment group were dominated by *Streptococcus*, which was annotated as susceptible to doxycycline in our database (Additional file 6). On days 21 and 28, the community shifted to a different low-diversity configuration where *Enterobacteriaceae*, annotated as resistant to doxycycline, was the most abundant taxon. Such radical and unanticipated shifts in community composition are worthy of further analysis with a microbiome response index, as we demonstrate in the next section.

### Extending microbiome data analysis with microbiome response indexes

Having demonstrated our antibiotic-specific microbiome response indexes across five studies, we wished to see how this new method could be used to extend existing methods of microbiome data analysis. We focused on associating MiRIx values with two summary measures of the microbiota: diversity and dysbiosis.

Antibiotic treatment has been shown to decrease microbiome diversity^7,8^, which is expected based on the intended effect of the drugs. However, inclusion of a microbiome response index allows for a more detailed analysis of the extent to which a decrease in diversity arises from a decrease in susceptible bacteria. For example, in the dataset from Basolo et al., we observed a positive correlation between the antibiotic-MiRIx and Shannon diversity (P=0.03), with vancomycin-treated subjects appearing low on both measures (Figure 4A). This result is consistent with elimination of susceptible bacterial species by the antibiotic: low-diversity states correspond to states where the response index has decreased following antibiotic exposure.

**Figure 4.**
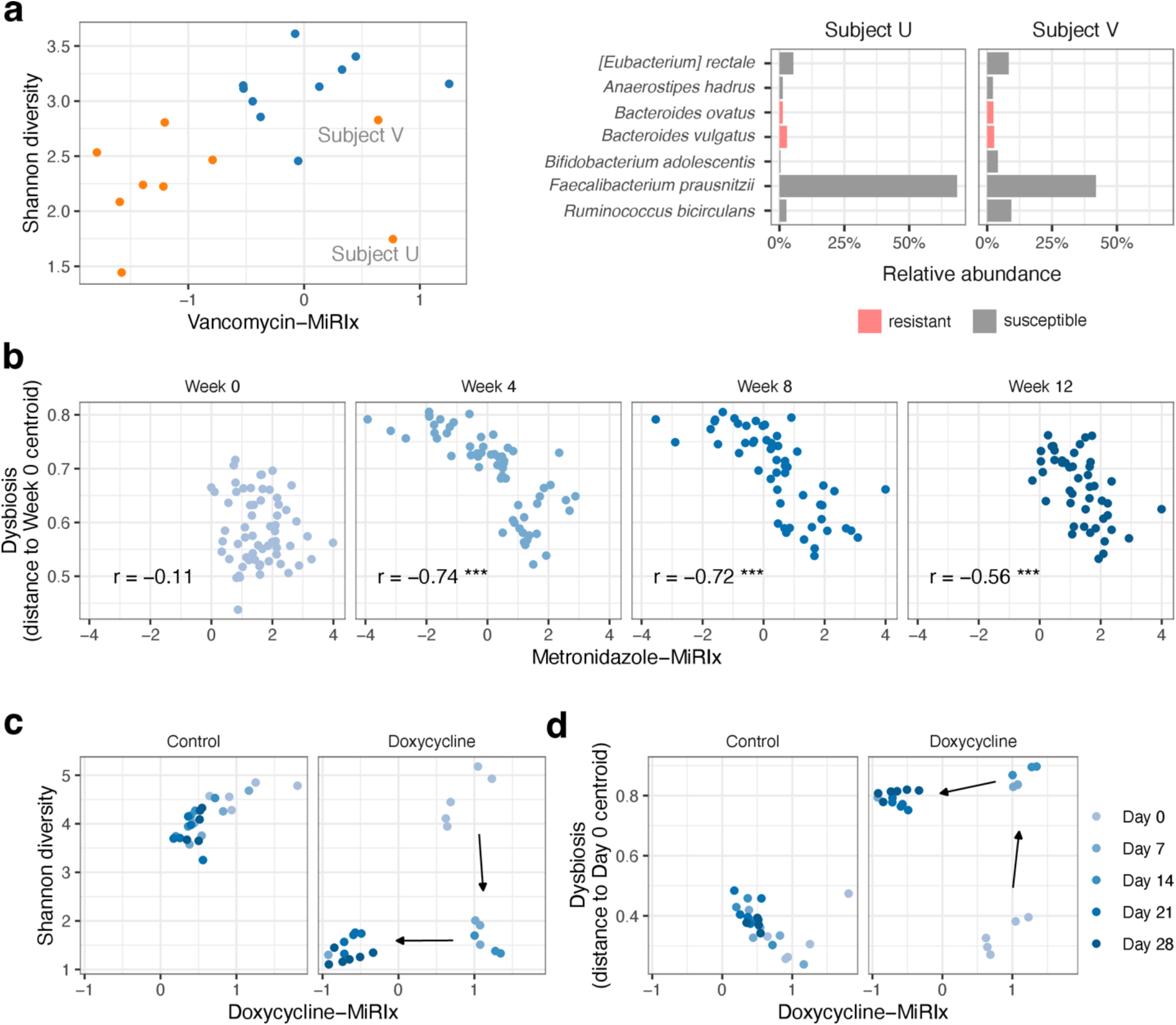
Extending microbiome data analysis with microbiome response indexes. (a) Vancomycin-MiRIx vs. Shannon diversity for the study from Boynton et al. A positive correlation is expected if vancomycin reduces the abundance of susceptible taxa. Taxonomic abundances are displayed for two vancomycin-treated subjects where the vancomycin-MiRIx did not decrease below the level in placebo controls. (b) Metronidazole-MiRIx vs. dysbiosis for the study from Sprockett et al. A negative correlation is expected if antibiotics increase the distance to healthy control samples by reducing the abundance of susceptible bacteria. (c) Doxycycline-MiRIx vs. Shannon diversity and (d) dysbiosis for the study from Boynton et al.

However, we noticed that two samples from vancomycin-treated subjects had a positive value of the vancomycin-MiRIx, indicating that most organisms were annotated as susceptible to the drug. One of the samples, from Subject U, had low Shannon diversity, similar to other antibiotic-treated subjects. The other sample, from Subject V, had higher diversity, in line with samples from the placebo group. We plotted the most abundant taxa and colored by vancomycin susceptibility. *Faecalibacterium prausnitzii*, annotated as vancomycin-susceptible in our database, was the most abundant species in both samples. In Subject U, *F. prausnitzii* accounted for over 60% of the relative abundance, corresponding to a low value for the Shannon diversity. In this way, we were able to frame differences in diversity and taxonomic abundance in terms of antibiotic susceptibility, using the microbiome response index as a starting point.

Dysbiosis refers to a microbiome’s departure from configurations observed in healthy subjects^35^, though there is some debate about the term’s definition^36,37^. Here, we adopt a simple quantitative definition based on each sample’s distance from a set of healthy reference communities, following the approach used by the Integrative Human Microbiome Project^38^. In the metronidazole data set from Sprockett et al., we observed that dysbiosis increased on weeks 4 and 8, relative to week 0 (P=10^−16^). Among the samples on weeks 4 and 8, the degree of dysbiosis was tightly correlated with the metronidazole-MiRIx (P=10^−11^ and 10^−9^ respectively), suggesting that the dysbiosis manifested as a reduction in antibiotic-susceptible organisms (Figure 4B). On week 12, the value of the metronidazole-MiRIx returned to baseline levels, but the dysbiosis remained elevated relative to baseline (P=10^−11^) and remained correlated with the metronidazole-MiRIx (P=0.0002). Thus, we were able to characterize the residual effects of antibiotic intervention, even after the balance of susceptible-vs-resistant organisms returned to baseline levels.

The doxycycline study from Boynton et al. provides an opportunity to relate diversity and dysbiosis to antibiotic response during a process of ecological succession. In the treatment group, the diversity decreased from baseline to day 7 (P=10^−11^), but there was no corresponding decrease in the doxycycline-MiRIx (P=0.3, Figure 4C). On day 21, the doxycycline-MiRIx decreased as the community adopted a different low-diversity configuration. Likewise, dysbiosis increased on day 7 (P=10^−14^) and was elevated through the study, but when plotted against the doxycycline-MiRIx, the two-stage succession process is readily seen (Figure 4D). Thus, we show how the microbiome response index can help to distinguish between stages of microbiome response to antibiotics and provide a context to the response that is relevant for the antibiotic used in the study.

In summary, we demonstrated that antibiotic MiRIx values can be integrated into the framework of existing microbiome data analysis methods, providing additional insight into how antibiotic exposure shapes microbial communities.

### Predicting antibiotic response in the healthy human microbiota

MiRIx values both quantify the state of the microbiome and give a qualitative prediction about how various antibiotics will impact a bacterial community. We applied the antibiotic-MiRIx approach to samples from various body sites in the human microbiome, hoping to gain insight about the susceptibility of these bacterial communities to various antibiotics in healthy adults. We downloaded data from the Human Microbiome Project (HMP) comprising three body sites—gut, oral, and skin—and computed MiRIx values for the five antibiotics presented previously in this paper: vancomycin, metronidazole, ciprofloxacin, amoxicillin, and doxycycline.

The MiRIx values for bacterial communities from the gut, oral cavity, and skin of healthy humans are shown in Figure 5A. For vancomycin, bacterial communities were roughly balanced between susceptible and non-susceptible organisms, with gut and oral communities tilted towards non-susceptible organisms and skin communities leaning towards susceptible organisms. For metronidazole and ciprofloxacin, targeting anaerobic and aerobic organisms respectively, bacterial communities from the anaerobic environment of the gut took on extreme values. For the broad-spectrum antibiotics amoxicillin and doxycycline, we observed that susceptible organisms outnumbered non-susceptible organisms by factors of 100-1000 to one. Moreover, we were unable to annotate any organisms as non-susceptible to amoxicillin or doxycycline in 27-36% of oral and skin microbiome samples.

**Figure 5.**
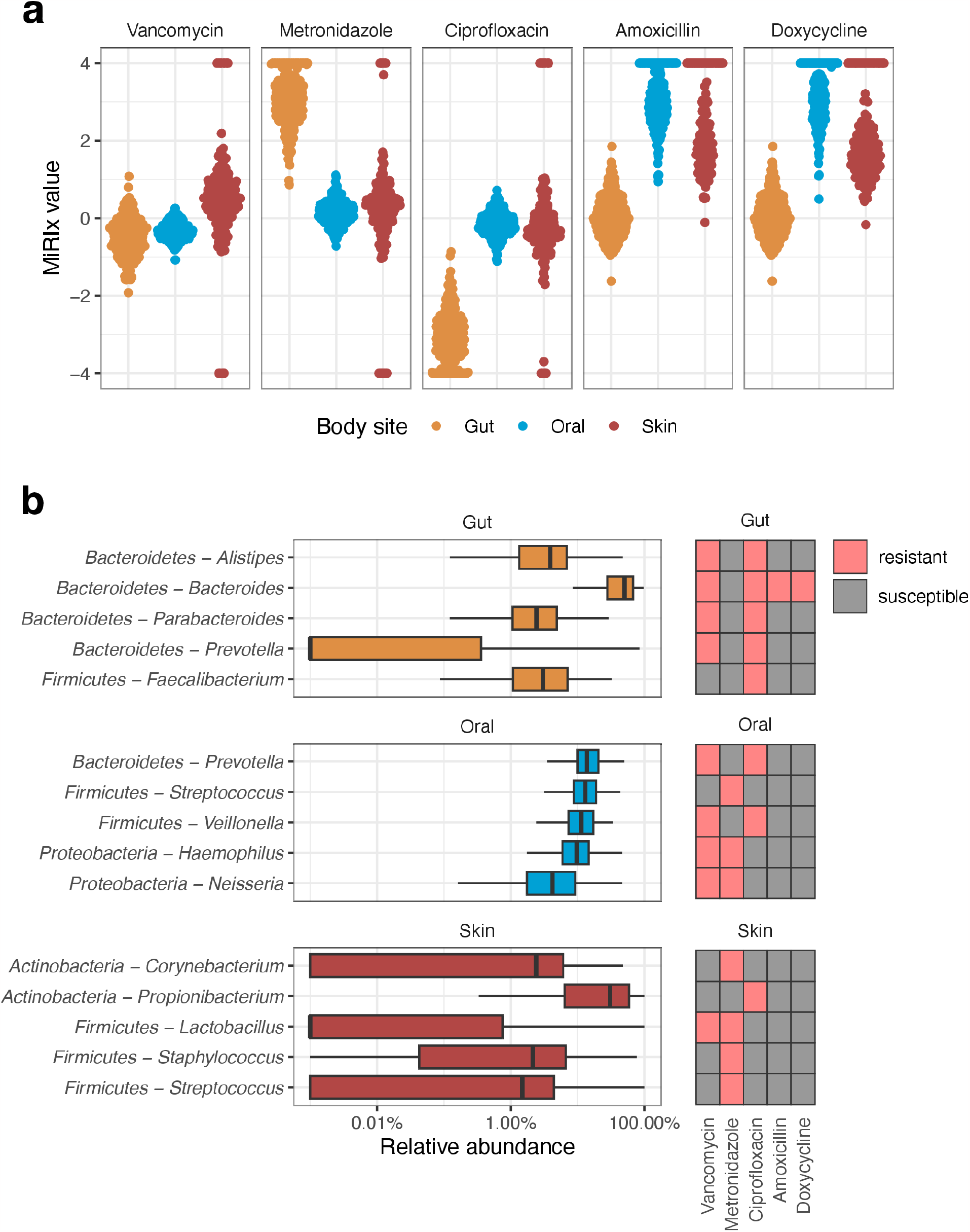
Predicting antibiotic response in the healthy human microbiota. (a) Antibiotic MiRIx values for samples from the Human Microbiome Project. (b) Taxonomic abundances and susceptibility for the top five taxa in each body site.

To investigate further, we selected the top five bacterial taxa at each body site and noted their annotations for susceptibility to each antibiotic (Figure 5B). In the gut, Gram-negative *Bacteroidetes* taxa were annotated as resistant and Gram-positive *Firmicutes* were annotated as susceptible to vancomycin, with *Veilonella* as a notable exception. Thus, the distribution of vancomycin-MiRIx values roughly reflects the balance of *Bacteroidetes* and *Firmicutes* in the human gut. For the oral samples, four of the five top taxa were annotated as resistant to vancomycin, with only *Streptococcus* annotated as susceptible. However, vancomycin-MiRIx values were narrowly distributed between -0.2 and -0.5 due to low overall variation of taxonomic abundances in the oral samples. In skin samples, four of the top five taxa were annotated as susceptible to vancomycin, with *Lactobacillus* as the exception.

The top five taxa in the gut were obligately anaerobic and were annotated as susceptible to metronidazole and non-susceptible to ciprofloxacin, corresponding to extremely high and low vancomycin-MiRIx values, respectively. The top oral taxa were balanced between obligate anaerobes and facultative anaerobes, though the MiRIx values for metronidazole and ciprofloxacin indicated a slight preference for obligate anaerobes like Prevotella and Veillonella. In the skin, only one genus in the top five was annotated as anaerobic and thus susceptible to metronidazole: *Propionibacterium*. However, the metronidazole-MiRIx and ciprofloxacin-MiRIx measures took on a wide range of values from skin samples, due to a high degree of variation in taxonomic abundance. In a small number of skin microbiome samples, we detected either no anaerobic or all anaerobic organisms.

We annotated all of the top five taxa in oral and skin samples as susceptible to the broad-spectrum antibiotics amoxicillin and doxycycline. In gut samples, we annotated *Bacteroides* as non-susceptible to both antibiotics. Because *Bacteroides* is a major component of the gut microbiome in the human population sampled by the HMP dataset, MiRIX values for both antibiotics were distributed between -0.3 and 0.4 in gut samples.

Although future research is needed to develop a more accurate picture of antibiotic impact on the human microbiome across body sites, we have demonstrated that our approach can provide a quantitative framework for studying antibiotic response in the microbiome when the data arrives. Our approach can be used to generate quantitative, sometimes surprising predictions, using basic rules applied from a standard underlying database. Additionally, we are able to point out specific bacterial taxa as the most important contributors to the antibiotic response.

## Discussion

Here, we introduced a new analytical approach, the Microbiome Response Index (MiRIx), which quantifies the balance of antibiotic-susceptible vs. antibiotic-resistant bacteria in a microbiome sample. The index values are calculated based on a database of bacterial phenotypes and antibiotic-specific susceptibilities, which were obtained from the literature and are distributed with our software. In addition to quantifying the state of the microbiome, the MiRIx values can be used to extend microbiome data analysis and make predictions about the state of the microbiome after hypothetical exposure to antibiotics.

We demonstrated our approach using five previously published data sets. Each time, we were able to derive new insight into microbiome composition and dynamics. Our approach revealed which bacterial taxa were responding in the expected manner, and which were responding in an unexpected way. In this sense, we find that our method is still useful even when it applies annotations that don’t reflect observations, because it allows researchers to pinpoint types of bacteria that defy our sometimes-naive expectations.

Given the appropriate database, our index is simple to calculate and understand: it is simply the balance of susceptible vs. resistant bacteria. However, many new analytical techniques can be crafted using this basic tool. MiRIx values can be combined with other microbiome measures to gain insight on how alpha and beta diversity change with antibiotic exposure. Values of the index provide a rough prediction of antibiotic response. Similarly, MiRIx values could also be correlated with clinical parameters such as route, dose and frequency of antibiotic administration to discern differences in microbiome response. With validation across clinically relevant antibiotic regimens the index in the future could be used to choose regimens with least microbiome disruption.

Our approach has several limitations and potential pitfalls. Because our approach uses a database to annotate bacterial taxa as resistant or susceptible to an antibiotic, its accuracy depends critically on the accuracy of the underlying database. We regard the state of the existing phenotype and resistance database as capturing broad aspects of antibiotic susceptibility, as described in medical textbooks and review articles. Although we argue that a quantitative index capturing the broad aspects of antibiotic susceptibility in the microbiome still delivers considerable value to researchers, we also wish for our method to be as accurate as possible. Extending and refining the database will be a target for future development; our database has substantial room to grow and capture more of the nuances in bacterial response to antibiotics.

Among such nuances are data-driven approaches that might extend the central MiRIx idea presented here. Firstly, because antibiotic susceptibility is often not all-or-nothing for bacterial species, it might be profitable to extend our approach with a probability-based estimation of antibiotic susceptibility for each taxon. Existing databases of antibiotic susceptibility rates, published by clinical laboratories, could be of great use in extending our approach in this way. Secondly, different bacterial strains may acquire antibiotic resistance through genes or mutations which can be detected in shotgun metagenomic sequencing. Our approach would benefit greatly from incorporation of sample-derived antibiotic resistance gene information. Both proposed extensions would introduce new conceptual complexity but would likely increase the accuracy of our approach and increase the number of connections to other relevant data sources.

## Conclusions

Here, we presented a new approach to quantify the balance of bacteria in a microbial community that are susceptible vs. resistant to a particular antibiotic. Based on evaluation of five previously published data sets, we demonstrated that our approach can be used to glean new, relevant insights from microbiome data analysis.

## Declarations

### Ethics approval and consent to participate

Not applicable.

### Consent for publication

Not applicable.

### Availability of data and materials

Software for computing microbiome response index (MiRIx) values is available from https://github.com/PennChopMicrobiomeProgram/mirix.

Code and data used to generate results presented in the manuscript are available from https://github.com/kylebittinger/mirix-paper.

### Authors’ contributions

Y.R. and K.B. designed the approach. V.T. and K.B. wrote the software. V.T., C.T., S.G.D., and K.B. carried out data analysis. S.M., H.L., and K.B. supervised the research and provided key guidance for the data analysis. V.T. and K.B. wrote the manuscript. All authors read and approved the final manuscript.

## Acknowledgements

Not applicable.

## Additional Files

**Additional file 1 (CSV format). Analysis of taxonomic abundance in the study from Basolo et al**. Taxa with a mean relative abundance greater than 0.005 were tested using linear models. The *estimate* column gives the estimated difference in log-transformed relative abundance (base 10) relative to the placebo group. The *fdr* column gives the p-value after adjustment to control for the false discovery rate.

**Additional file 2 (CSV format). Analysis of taxonomic abundance in the study from Sprockett et al**. Taxa with a mean relative abundance greater than 0.005 were tested using linear mixed effects models. The *term* column gives the week of the study where abundances were compared. The *estimate* column gives the estimated difference in log-transformed relative abundance (base 10) relative to the abundance at week 0. The *fdr* column gives the p-value after adjustment to control for the false discovery rate.

**Additional file 3 (CSV format). Analysis of taxonomic abundance in the study from Willmann et al**. Taxa with a mean relative abundance greater than 0.005 were tested using linear mixed effects models. The *term* column gives the day of the study where abundances were compared. The *estimate* column gives the estimated difference in log-transformed relative abundance (base 10) relative to the abundance at day 0. The *fdr* column gives the p-value after adjustment to control for the false discovery rate.

**Additional file 4 (PDF format). Relative abundance of antibiotic resistance genes in the study from Willmann et al**. Genes conferring resistance to fluoroquinolone antibiotics with a mean relative abundance of at least 10^−5^ are shown. Stars indicate a statistically significant difference relative to the abundance on day 0, after correction for multiple comparisons (*** P<0.001, ** P<0.01, * P<0.05).

**Additional file 5 (CSV format). Analysis of taxonomic abundance in the study from Cabral et al**. Taxa with a mean relative abundance greater than 0.005 were tested using linear models. The *estimate* column gives the estimated difference in log-transformed relative abundance (base 10) relative to the control group. The *fdr* column gives the p-value after adjustment to control for the false discovery rate.

**Additional file 6 (CSV format). Analysis of taxonomic abundance in the study from Boynton et al**. Taxa with a mean relative abundance greater than 0.005 were tested using linear mixed effects models. The *term* column gives the day of the study where abundances were compared. The *estimate* column gives the estimated difference in log-transformed relative abundance (base 10) relative to the abundance at day 0. The *fdr* column gives the p-value after adjustment to control for the false discovery rate.

## References

1 Maier, L. et al. Extensive impact of non-antibiotic drugs on human gut bacteria. Nature 555, 623–628 (2018). 10.1038/nature25979

2 Bhalodi, A. A., van Engelen, T. S. R., Virk, H. S. & Wiersinga, W. J. Impact of antimicrobial therapy on the gut microbiome. J Antimicrob Chemother 74, i6–i15 (2019). 10.1093/jac/dky530

3 Zimmermann, P. & Curtis, N. The effect of antibiotics on the composition of the intestinal microbiota - a systematic review. J Infect 79, 471–489 (2019). 10.1016/j.jinf.2019.10.008

4 Palleja, A. et al. Recovery of gut microbiota of healthy adults following antibiotic exposure. Nat Microbiol 3, 1255–1265 (2018). 10.1038/s41564-018-0257-9

5 Tanes, C. et al. Role of dietary fiber in the recovery of the human gut microbiome and its metabolome. Cell Host Microbe 29, 394–407 e395 (2021). 10.1016/j.chom.2020.12.012

6 Lewis, J. D. et al. Inflammation, Antibiotics, and Diet as Environmental Stressors of the Gut Microbiome in Pediatric Crohn’s Disease. Cell Host Microbe 18, 489–500 (2015). 10.1016/j.chom.2015.09.008

7 Dethlefsen, L., Huse, S., Sogin, M. L. & Relman, D. A. The pervasive effects of an antibiotic on the human gut microbiota, as revealed by deep 16S rRNA sequencing. PLoS Biol 6, e280 (2008). 10.1371/journal.pbio.0060280

8 Dethlefsen, L. & Relman, D. A. Incomplete recovery and individualized responses of the human distal gut microbiota to repeated antibiotic perturbation. Proc Natl Acad Sci U S A 108 Suppl 1, 4554–4561 (2011). 10.1073/pnas.1000087107

9 Klein, E. Y., Tseng, K. K., Pant, S. & Laxminarayan, R. Tracking global trends in the effectiveness of antibiotic therapy using the Drug Resistance Index. BMJ Glob Health 4, e001315 (2019). 10.1136/bmjgh-2018-001315

10 Laxminarayan, R. & Klugman, K. P. Communicating trends in resistance using a drug resistance index. BMJ Open 1, e000135 (2011). 10.1136/bmjopen-2011-000135

11 Ciccolini, M., Spoorenberg, V., Geerlings, S. E., Prins, J. M. & Grundmann, H. Using an index-based approach to assess the population-level appropriateness of empirical antibiotic therapy. J Antimicrob Chemother 70, 286–293 (2015). 10.1093/jac/dku336

12 Chitanand, M. P., Kadam, T. A., Gyananath, G., Totewad, N. D. & Balhal, D. K. Multiple antibiotic resistance indexing of coliforms to identify high risk contamination sites in aquatic environment. Indian J Microbiol 50, 216–220 (2010). 10.1007/s12088-010-0042-9

13 Hughes, J. S. et al. How to measure the impacts of antibiotic resistance and antibiotic development on empiric therapy: new composite indices. BMJ Open 6, e012040 (2016). 10.1136/bmjopen-2016-012040

14 Port, J. A., Cullen, A. C., Wallace, J. C., Smith, M. N. & Faustman, E. M. Metagenomic frameworks for monitoring antibiotic resistance in aquatic environments. Environ Health Perspect 122, 222–228 (2014). 10.1289/ehp.1307009

15 Li, X., Zhang, Z., Liang, B., Ye, F. & Gong, W. A review: antimicrobial resistance data mining models and prediction methods study for pathogenic bacteria. J Antibiot (Tokyo) 74, 838–849 (2021). 10.1038/s41429-021-00471-w

16 Basolo, A. et al. Effects of underfeeding and oral vancomycin on gut microbiome and nutrient absorption in humans. Nat Med 26, 589–598 (2020). 10.1038/s41591-020-0801-z

17 Willmann, M. et al. Distinct impact of antibiotics on the gut microbiome and resistome: a longitudinal multicenter cohort study. BMC Biol 17, 76 (2019). 10.1186/s12915-019-0692-y

18 Clarke, E. L. et al. Sunbeam: an extensible pipeline for analyzing metagenomic sequencing experiments. Microbiome 7, 46 (2019). 10.1186/s40168-019-0658-x

19 Martin, M. Cutadapt Removes Adapter Sequences From High-Throughput Sequencing Reads. EMBnet.journal 17, 10–12 (2011).

20 Bolger, A. M., Lohse, M. & Usadel, B. Trimmomatic: a flexible trimmer for Illumina sequence data. Bioinformatics 30, 2114–2120 (2014). 10.1093/bioinformatics/btu170

21 Li, H. & Durbin, R. Fast and accurate short read alignment with Burrows-Wheeler transform. Bioinformatics 25, 1754–1760 (2009). 10.1093/bioinformatics/btp324

22 Wood, D. E., Lu, J. & Langmead, B. Improved metagenomic analysis with Kraken Genome Biol 20, 257 (2019). 10.1186/s13059-019-1891-0

23 Sprockett, D. et al. Treatment-Specific Composition of the Gut Microbiota Is Associated With Disease Remission in a Pediatric Crohn’s Disease Cohort. Inflamm Bowel Dis 25, 1927–1938 (2019). 10.1093/ibd/izz130

24 Boynton, F. D. D., Ericsson, A. C., Uchihashi, M., Dunbar, M. L. & Wilkinson, J. E. Doxycycline induces dysbiosis in female C57BL/6NCrl mice. BMC Res Notes 10, 644 (2017). 10.1186/s13104-017-2960-7

25 Bolyen, E. et al. Reproducible, interactive, scalable and extensible microbiome data science using QIIME 2. Nat Biotechnol 37, 852–857 (2019). 10.1038/s41587-019-0209-9

26 Callahan, B. J. et al. DADA2: High-resolution sample inference from Illumina amplicon data. Nat Methods 13, 581–583 (2016). 10.1038/nmeth.3869

27 Bokulich, N. A. et al. Optimizing taxonomic classification of marker-gene amplicon sequences with QIIME 2’s q2-feature-classifier plugin. Microbiome 6, 90 (2018). 10.1186/s40168-018-0470-z

28 Schiffer, L. et al. HMP16SData: Efficient Access to the Human Microbiome Project Through Bioconductor. Am J Epidemiol 188, 1023–1026 (2019). 10.1093/aje/kwz006

29 Benjamini, Y. & Hochberg, Y. Controlling the false discovery rate: a practical and powerful approach to multiple testing. Journal of the Royal Statistical Society, Series B 57, 289–300 (1995).

30 Anderson, M. J. A new method for non-parametric multivariate analysis of variance. Austral Ecology 26, 32–46 (2001).

31 Edlund, C., Barkholt, L., Olsson-Liljequist, B. & Nord, C. E. Effect of vancomycin on intestinal flora of patients who previously received antimicrobial therapy. Clin Infect Dis 25, 729–732 (1997). 10.1086/513755

32 Waters, J. L. & Salyers, A. A. Regulation of CTnDOT conjugative transfer is a complex and highly coordinated series of events. mBio 4, e00569–00513 (2013). 10.1128/mBio.00569-13

33 Cabral, D. J. et al. Microbial Metabolism Modulates Antibiotic Susceptibility within the Murine Gut Microbiome. Cell Metab 30, 800–823 e807 (2019). 10.1016/j.cmet.2019.08.020

34 Reygaert, W. C. An overview of the antimicrobial resistance mechanisms of bacteria. AIMS Microbiol 4, 482–501 (2018). 10.3934/microbiol.2018.3.482

35 Petersen, C. & Round, J. L. Defining dysbiosis and its influence on host immunity and disease. Cell Microbiol 16, 1024–1033 (2014). 10.1111/cmi.12308

36 Hooks, K. B. & O’Malley, M. A. Dysbiosis and Its Discontents. mBio 8 (2017). 10.1128/mBio.01492-17

37 Brussow, H. Problems with the concept of gut microbiota dysbiosis. Microb Biotechnol 13, 423–434 (2020). 10.1111/1751-7915.13479

38 Lloyd-Price, J. et al. Multi-omics of the gut microbial ecosystem in inflammatory bowel diseases. Nature 569, 655–662 (2019). 10.1038/s41586-019-1237-9

