## Supplementary figures and images for "A quantitative approach to measure and predict microbiome response to antibiotics"

### Additional file 4

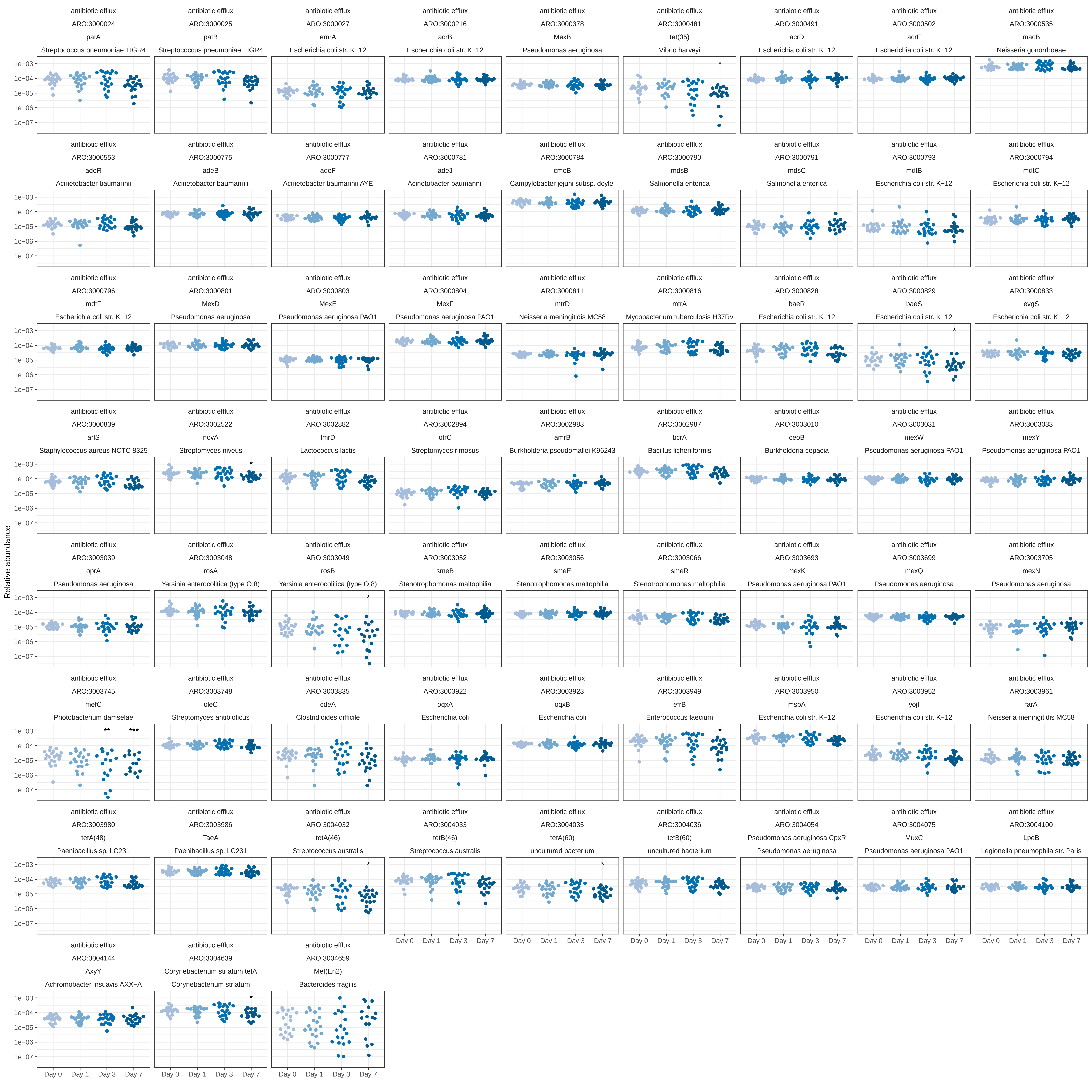
